# Bidirectional synaptic changes in deep and superficial hippocampal neurons following *in vivo* activity

**DOI:** 10.1101/2022.10.27.514077

**Authors:** Marcus Berndt, Massimo Trusel, Todd F. Roberts, Lenora J. Volk, Brad E. Pfeiffer

## Abstract

Neuronal activity during experience is thought to induce plastic changes within the hippocampal network which underlie memory formation, although the extent and details of such changes *in vivo* remain unclear. Here, we employed a temporally precise marker of neuronal activity, CaMPARI2, to label active CA1 hippocampal neurons *in vivo*, followed by immediate acute slice preparation and electrophysiological quantification of synaptic properties. Recently active neurons in the superficial sublayer of *stratum pyramidale* displayed larger post-synaptic responses at excitatory synapses from area CA3, with no change in pre-synaptic release probability. In contrast, *in vivo* activity weakened both pre- and post-synaptic excitatory weights onto pyramidal cells in the deep sublayer. *In vivo* activity of deep and superficial neurons within sharp-wave ripples was bidirectionally changed across experience, consistent with the observed changes in synaptic weights. These findings reveal novel, fundamental mechanisms through which the hippocampal network is modified by experience to store information.

## INTRODUCTION

Considerable evidence indicates that storage of information in the brain requires synaptic plasticity^1–4^, but the precise synaptic changes induced by experience are not well understood. Previous strategies to quantify experience-driven changes in *in vivo* synaptic function have been limited in their ability to quantify cell-type or input specific synaptic changes^5–8^ or require hours-long timescales to label active populations *in vivo*^9,10^. While prior methods have produced considerable insight into neural mechanisms of memory and plasticity, the nature and extent of modifications to the intact hippocampal network driven by natural experience remain unclear. CaMPARI2 (calcium-modulated photo-activatable ratiometric integrator, 2^nd^ generation)^11,12^ is an engineered green fluorescent protein with native excitation/emission of 502/516 nm. When bound to calcium, near-UV light stimulation produces a covalent break, permanently converting CaMPARI2 to 562/577 nm excitation/emission^11,12^. Unlike immediate early gene expression-based labeling strategies, which are not directly correlated to neural activity rates^13,14^, CaMPARI2 photoconversion is dependent upon intracellular calcium levels^11^ and linearly correlated to neural activity^12^, thus making it a powerful experimental method to identify active vs. inactive neurons on a sub-second timescale to query the synaptic consequences of *in vivo* activity patterns.

Experience-dependent plasticity across the hippocampal formation is critical for the formation of episodic and spatial memories^15–17^. During navigation, excitatory neurons in the hippocampus, termed “place cells,” are selectively active in restricted spatial regions, forming a spatially and temporally organized representation of experience across the hippocampal network^18,19^. Establishment and maintenance of this cognitive map is heavily dependent upon synaptic plasticity at individual hippocampal neurons^5–10,20–22^, but it is unclear how such changes might impact subsequent memory retrieval. In a well-established example of network-level memory expression, patterns of hippocampal population activity that occur during behavior can be reactivated during off-line periods such as quiet rest or slow-wave sleep^23–25^. Off-line reactivation, often referred to as “replay,” is temporally compressed and occurs within local field potential (LFP) events termed sharp-wave/ripples (SWRs)^26,27^. While pre-existing network connectivity patterns may influence the content of hippocampal SWRs^28–31^, high-fidelity expression of post-experience replay events is dependent on NMDA receptor function during exploration^32–34^, arguing that synaptic plasticity occurring during behavior modifies the hippocampal network to support subsequent reactivation, although the nature of these changes is unknown.

It is increasingly evident that the excitatory neurons of the hippocampal network do not form a homogenous population, but rather can be differentiated into at least two distinct categories based on development^31,35^, gene expression^36,37^, and synaptic properties^38,39^. These two populations are largely segregated along the radial axis of *stratum pyramidale* and are canonically termed the deep layer (nearest *stratum oriens*) and the superficial layer (nearest *stratum radiatum*)^40^. Deep and superficial neurons have divergent local and interregional synaptic organization^31,39,41–43^, which may underlie differences in place cell properties and the stability of spatial representations between these two populations^31,38,44–49^. However, it is unknown whether synapses onto deep and superficial neurons are differentially impacted by *in vivo* activity or how such changes may impact the ability of the hippocampal network to store and retrieve information.

## RESULTS

### CaMPARI2 Labels Active Hippocampal Populations *In Vivo*

To study the synaptic consequences of *in vivo* activity within the hippocampal network, we expressed CaMPARI2 in hippocampal dorsal area CA1 of adult rats. CaMPARI2 has native green fluorescence but is permanently converted to red fluorescence in the combined presence of high intracellular calcium and near-UV light (Figure 1 A)^11,12^. While individual CaMPARI2 molecules are binary (existing in either the green or red fluorescent state), the fluorescence of the entire population of CaMPARI2 within a given cell provides an analog, ratiometric measure of overall activity levels of that neuron during photostimulation^11,12^. Following viral expression of CaMPARI2 in excitatory hippocampal neurons, we implanted optic fibers above dorsal area CA1 to label active CA1 pyramidal neurons with sub-second temporal precision via CaMPARI2 photoconversion (Figure 1 B). We allowed rats to freely explore a novel, rewarded linear track and provided 405 nm light stimulation during periods of active movement, permanently photoconverting CaMPARI2 from green to red in highly active neurons (*i.e*., place cells with place fields in this novel environment). In a control cohort, we provided equivalent duration and intensity of photostimulation while rats quietly rested in their familiar home cage. For both cohorts, rats were sacrificed immediately following photostimulation and acute hippocampal slices were rapidly prepared for live tissue imaging and electrophysiological analysis. Photoconversion was restricted to a cone beneath the optic fiber (Figure 1 C), confirming that photoconversion was governed by light presentation.

**Figure 1.**
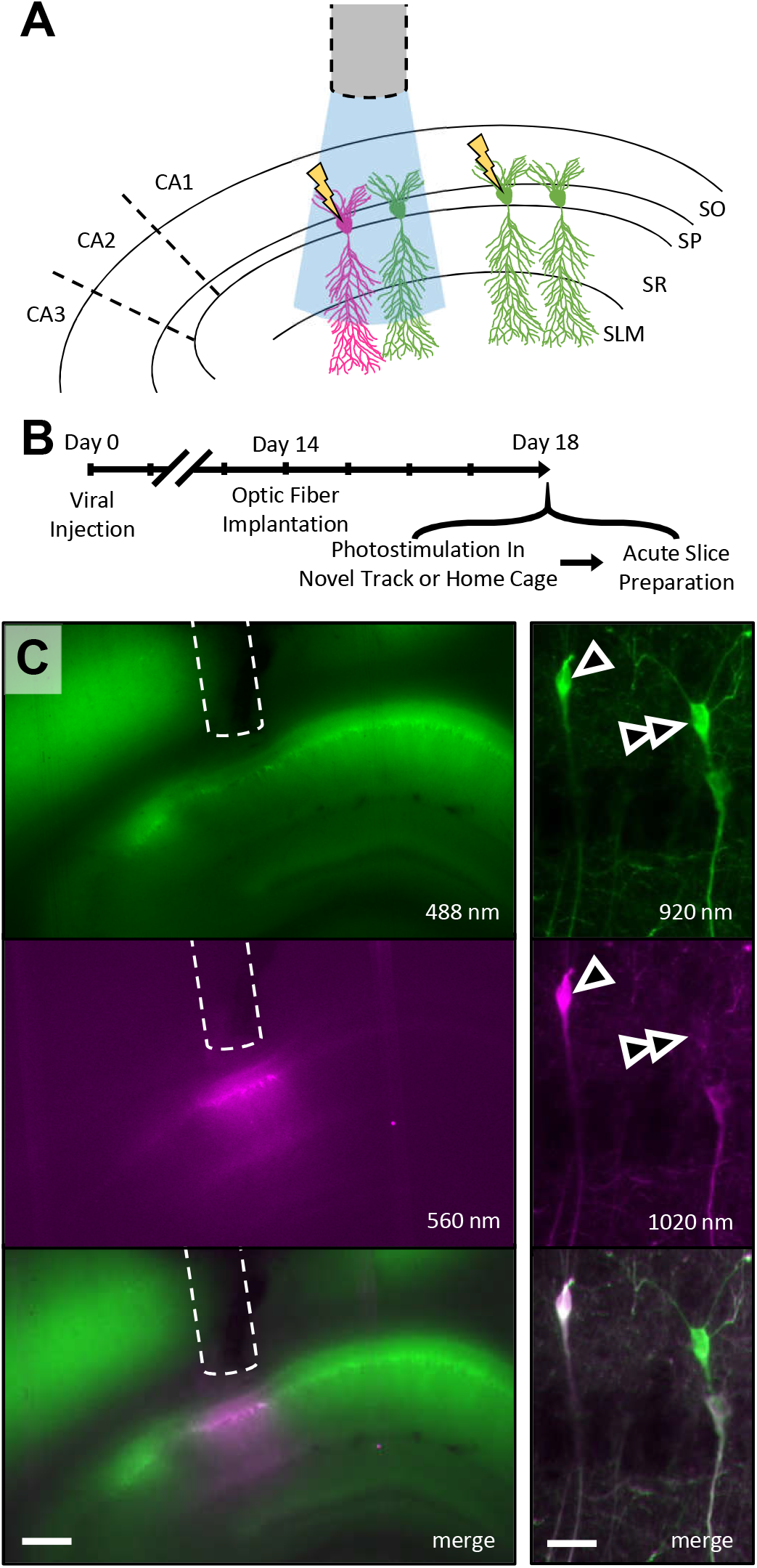
Experiment design and CaMPARI2 expression and conversion. **A.** Schema of CaMPARI2 experiment design. CaMPARI2-expressing CA1 pyramidal neurons initially display green fluorescence. In the combined presence of high intracellular calcium produced by neural activity (lightning bolt) and 405 nm light provided by the implanted optic fiber (blue cone), CaMPARI2 is permanently converted to red fluorescence, persistently marking neurons that were active within the optic cone during light presentation. SO, *stratum oriens*; SP, *stratum pyramidale*; SR, *stratum radiatum*; SLM, *stratum lacunosum/moleculare*. **B**. Timeline of experiment. **C**. Left, widefield CaMPARI2 fluorescence in acute dorsal hippocampal slice following photostimulation. Top, fluorescence of native (unconverted), green CaMPARI2. Middle, fluorescence of permanently photoconverted, red CaMPARI2. Bottom, merged of top and middle. Optic fiber tract denoted by dashed line. Scale bar, 400 μm; excitation wavelength indicated. Right, as left, for mean projection two-photon z-stack image of CA1 pyramidal layer showing both native and photoconverted CaMPARI2-expressing neurons. Arrowhead denotes a strongly photoconverted neuron; double-arrowhead denotes unconverted neuron. Scalebar, 40 μm.

Via two-photon imaging of live acute slices, we quantified the ratio of red-to-green fluorescence in each CaMPARI2-expressing pyramidal neuron within the optic cone as a measure of activity level during photostimulation (Figure 2 A-B). In the experimental cohort, in which neurons were photostimulated during active exploration, we observed a bimodal distribution of photoconversion ratios (Figure 2 A), consistent with a population of highly active (photoconverted) place cells and a population of silent (unconverted) cells^50^. We conservatively defined neurons with a red/green ratio lower than 0.4 as unconverted and neurons with a ratio of 0.6 or higher as photoconverted (Figure 2 A). To provide confidence to our classifications, neurons with conversion ratios between 0.4 and 0.6 were unclassified and not included in subsequent analyses. With these criteria, 55 of 147 (37.4%) CaMPARI2-expressing neurons beneath the optic fiber were photoconverted during active exploration, consistent with reports of the relative ratios of active place cells and silent cells^50,51^. In contrast, we observed a much smaller percentage of photoconverted neurons (10 of 132, 7.6%) in rats which received photo-stimulation during home cage quiet rest (Figure 2 B), consistent with reduced overall levels of activity during immobility in familiar environments^9^. The percentage of neurons that were converted during active exploration was significantly higher than the percentage converted during quiet rest (binomial cumulative distribution, *p* < 10^−10^). To assess the degree of photoconversion within individual neurons, we directly compared red/green ratios for converted and unconverted neurons in both cohorts, as well as for neurons outside of the photoconversion cone (Figure 2 C). Photoconverted neurons did not significantly differ in their red/green ratio between the active exploration and home cage cohorts (Figure 2 C), indicating that while the home cage cohort had fewer active neurons, the overall activity level of that active population was roughly equivalent to the activity levels of place cells in the active exploration cohort, consistent with a small population of neurons with high basal activity rates in the home cage^52^. Furthermore, the red/green ratios of neurons that were classified as unconverted in either the home cage or active exploration cohorts were not significantly different from the red/green ratios of neurons that were outside of the optic light cone (Figure 2 C), supporting our classification of this population as “unconverted.”

**Figure 2.**
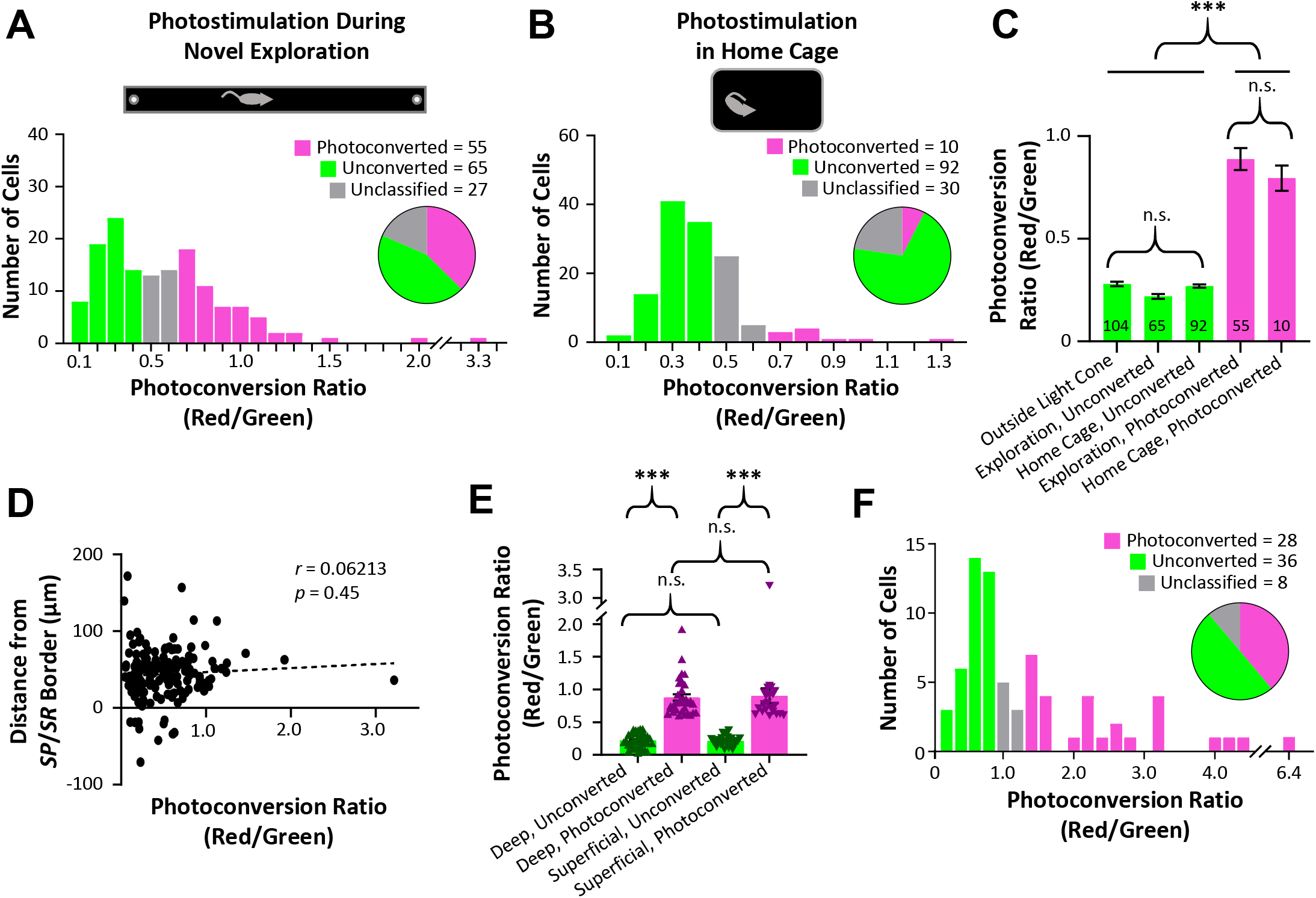
Distribution of CaMPARI2 photoconversion during novel exploration vs. home cage rest. **A**. Histogram of photoconversion ratios across all CaMPARI2-expressing pyramidal cells (*n* = 147 cells, 3 rats) within the optic cone photostimulated during active exploration of a novel linear track. **B**. As (A), for photostimulation during quiet rest in familiar home cage (*n* = 132 cells, 3 rats). **C**. Mean ± S.E.M. photoconversion ratio for neurons outside of the light cone (far left), and neurons classified as Unconverted or Photoconverted for both novel exploration and home cage cohorts. **D**. Plot of photoconversion ratio and distance from the *stratum pyramidale*/*stratum radiatum* border for neurons optically stimulated during novel exploration and imaged on two-photon microscope. **E**. Mean ± S.E.M. and individual data points of photoconversion ratio for neurons in (D) classified as deep vs. superficial and photoconverted vs. unconverted. **F**. Histogram of photoconversion ratios on electrophysiology microscope across CaMPARI2-expressing pyramidal cells used in whole cell electrophysiology experiments (*n* = 72 cells, 14 rats) within the optic cone photostimulated during active exploration of a novel linear track. Statistical tests and results: (C) One-way ANOVA (4,321)=138.2; *p* < 10^−4^. *Post hoc* multiple comparisons (Tukey HSD): Outside vs. Either Unconverted, *p* > 0.25; Exploration Photoconverted vs. Home Cage Photoconverted, *p* > 0.6; Outside/Either Unconverted vs. Either Photoconverted *p* < 10^−4^. (E) Two-way ANOVA Deep vs. Superficial F(1,116)=0.0257, *p* = 0.8730; Unconverted vs. Photoconverted F(1,116)=163.2, *p* < 10^−4^; Interaction F(1,116)=0.113, *p* = 0.7375. Multiple comparisons (Tukey HSD): Either-Unconverted vs. Either-Photoconverted *p* < 10^−4^; Deep-Unconverted vs. Superficial Unconverted *p* = 0.9992; Deep-Photoconverted vs. Superficial Photoconverted *p* = 0.987.

Given established differences in spatial encoding between deep and superficial pyramidal neurons^31,38,44–49^, we quantified the photoconversion ratio of each neuron as a function of somatic distance from the border between *stratum pyramidale* and *stratum radiatum*. There was no correlation between radial location and photoconversion ratio (Figure 2 D). We categorized superficial and deep neurons, respectively, as cells within or greater than 40 μm from the *stratum pyramidale*/*stratum radiatum* border^41^ and found no difference in photoconversion rates or red/green ratios between deep and superficial neurons (Figure 2 E, deep cells: 33 of 85 cells (38.8%) classified as photoconverted; superficial cells: 22 of 62 cells (35.5%) classified as photoconverted; binomial cumulative distribution, *p* = 0.22).

### Recent Activity Differentially Impacts Synaptic Strength onto Deep and Superficial Neurons

We next quantified the synaptic consequences of *in vivo* activity by performing whole cell electrophysiology on converted and unconverted neurons within the optic light cone from a second novel exploration cohort. As this experiment was performed with a separate fluorescence microscope on an electrophysiology rig (with different fluorescence sensitivity than the two-photon microscope used for the previous analyses), we re-assessed the distribution of red/green fluorescence ratios from this cohort. While overall red/green ratios were slightly higher on the electrophysiology rig microscope than on the two-photon microscope, the distribution of conversion ratios remained bimodal, with 38.9% (28 of 72) of CaMPARI2-expressing neurons displaying strong photoconversion (Figure 2 F), statistically similar to the rate of conversion identified under the two-photon microscope (Figure 2 A, binomial cumulative distribution, *p* = 0.35). We observed no difference in capacitance or input resistance between photoconverted and unconverted neurons [Input resistance mean ± SEM: deep converted (n = 14) 209.5 ± 29.97; deep unconverted (n = 10) 221.9 ± 53.94; superficial converted (n = 9) 207.1 ± 25.62; superficial unconverted (n = 11) 130.7 ± 20.04; two-way ANOVA deep vs. superficial F(1,40)=1.816, *p* = 0.1854; unconverted vs. photoconverted F(1,40)=0.849, *p* = 0.3624; interaction F(1,40)=1.634, *p* = 0.2085. Capacitance mean ± SEM: deep converted (n = 12) 64.76 ± 15.36; deep unconverted (n = 10) 73.66 ± 17.53; superficial converted (n = 9) 65.92 ± 27.25; superficial unconverted (n = 10) 104.9 ± 20.67; two-way ANOVA deep vs. superficial F(1,37)=0.6533, *p* = 0.4241; unconverted vs. photoconverted F(1,37)=1.427, *p* = 0.2399; interaction F(1,37)=0.5631, *p* = 0.4577].

We directly stimulated the Schaffer collaterals to measure excitatory CA3 input to unconverted or photoconverted CA1 neurons and used paired-pulse ratio (PPR) and the ratio of AMPA/NMDA responses to measure pre-synaptic release probability and post-synaptic strength, respectively. Deep pyramidal cells displayed a positive relationship between the CaMPARI2 red/green conversion ratio and the PPR of Schaffer collateral synapses (Figure 3 A). As higher PPR values are consistent with reduced pre-synaptic release probability, these data indicate that *in vivo* activity of deep pyramidal neurons depresses pre-synaptic excitatory inputs to these cells from area CA3. In addition, the ratio of AMPA/NMDA responses onto deep pyramidal neurons were negatively correlated with photoconversion ratios (Figure 3 B). These results collectively argue that recent neural activity during novel exploration significantly depresses both pre- and post-synaptic weights between excitatory CA3 neurons and deep CA1 pyramidal neurons. We next assessed CA3 synaptic strength onto superficial CA1 neurons (Figure 3 C-D). In contrast to what was observed for deep pyramidal neurons, PPR values onto superficial pyramidal neurons were not affected by *in vivo* activity (Figure 3 C). However, the post-synaptic weights of superficial neurons showed a positive correlation with the CaMPARI2 conversion ratio (Figure 3 D). These data suggest that *in vivo* activity during active exploration produces post-synaptic potentiation of Schaffer collateral synapses onto superficial neurons. To further quantify the effects of activity on synaptic function, we categorized deep and superficial neurons into photoconverted vs. unconverted populations. Photoconverted deep pyramidal neurons had a significantly higher PPR and a significantly lower AMPA/NMDA ratio than unconverted deep neurons, while photoconverted superficial neurons had a significantly higher AMPA/NMDA ratio than unconverted superficial neurons (Figure 3 E-F).

**Figure 3.**
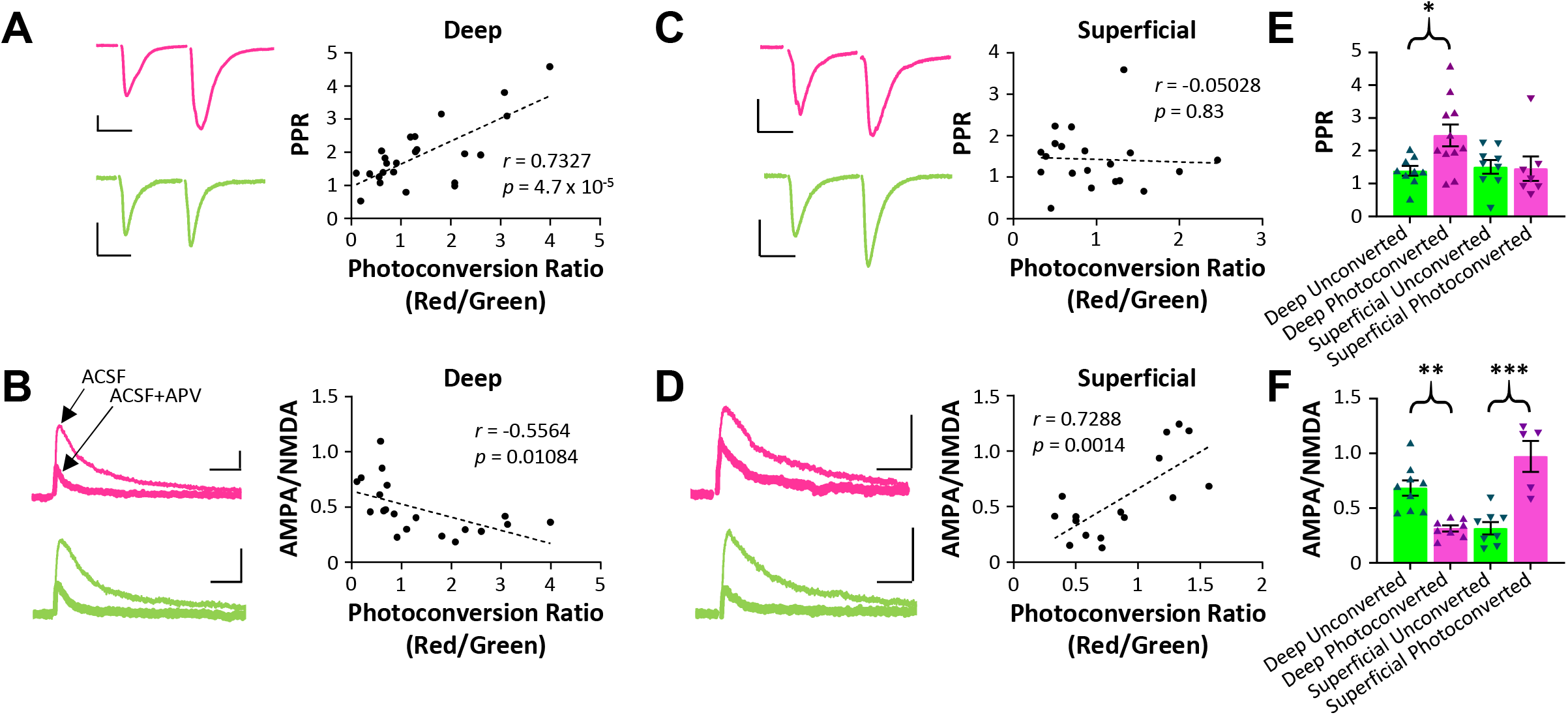
Deep and superficial cells display distinct synaptic differences between unconverted and photoconverted neurons. **A.** Left, representative paired-pulse traces from unconverted (green) and photoconverted (magenta) deep pyramidal neurons. Scale bar 100 pA, 25 ms. Right, PPR vs. photoconversion ratio for all deep neurons. **B**. Left, representative AMPA/NMDA traces from unconverted (green) and photoconverted (magenta) deep pyramidal neurons held at −40 mV. Thin trace is ACSF-only, thick trace is ACSF with 100 mM AP5. Scale bar 50 pA, 50 ms. Right, AMPA/NMDA ratio vs. photoconversion ratio for all deep neurons. **C-D**. As (A-B), for superficial pyramidal neurons. **E**. Mean ± S.E.M. and individual data points of PPR for neurons in (A) and (C) classified as photoconverted vs. unconverted. **F**. Mean ± S.E.M. and individual data points of AMPA/NMDA ratio for neurons in (B) and (D) classified as photoconverted vs. unconverted. Statistical tests and results: (E) Two-way ANOVA Deep vs. Superficial F(1,32)=2.453, *p* = 0.1271; Unconverted vs. Photoconverted F(1,32)=3.257, *p* = 0.0805; Interaction F(1,32)=3.942, *p* = 0.0557. Multiple comparisons (Tukey HSD): Unconverted-Deep vs. Photoconverted-Deep *p* = 0.0365. (F) Two-way ANOVA Deep vs. Superficial F(1,26)=3.986, *p* = 0.0565; Unconverted vs. Photoconverted F(1,26)=3.974, *p* = 0.0568; Interaction F(1,26)=50.13, *p* < 10^−4^. Multiple comparisons (Tukey HSD): Unconverted-Deep vs. Photoconverted-Deep *p* = 0.003; Unconverted-Superficial vs. Photoconverted-Superficial *p* < 10^−4^.

### Exploration Induces Bidirectional Effects on Deep and Superficial Activity in Off-Line Reactivation

Together, the above findings argue that *in vivo* activity produces distinct and input-specific synaptic changes to deep versus superficial CA1 neurons. Specifically, network activity during exploration appears to potentiate excitatory CA3 input to superficial place cells but depress input to deep place cells. During sharp-wave/ripples (SWRs), the dominant excitatory input onto CA1 neurons originates from CA3^53–55^, and the strength of CA3-to-CA1 synapses is positively correlated with participation of CA1 neurons in SWR events^56^. Thus, we hypothesized that *in vivo* activity during exploration should differentially impact the activity of deep and superficial neurons during SWRs following the experience^41^. To test this hypothesis, we obtained large-scale *in vivo* electrophysiological recordings from dorsal area CA1 in a separate cohort of freely behaving rats and analyzed neural activity within SWRs before and after exploration of a linear track (Figure 4 A). The sharp-wave component of the SWR reverses across the pyramidal layer^57^, allowing us to assess the radial depth of each tetrode via the sharp-wave deflection (Figure 4 B). In each recording session, we observed a large distribution of recording depths (Figure 4 C). To ensure our analysis was limited to deep vs. superficial sublayers, we eliminated any tetrode with a sharp-wave deflection between 50 and −50 μV (Figure 4 C). We further restricted our analyses to unit clusters with large amplitude spikes (mean amplitude >200 μV) to ensure that the cells producing those action potentials were located near the recording site, thus allowing us to identify deep neurons on tetrodes with upward-deflecting sharp-waves and superficial neurons on tetrodes with downward-deflecting sharp-waves. When we quantified the rates of participation of deep and superficial cells in pre-experience SWRs, we observed no significant difference between the two populations (Figure 4 D). However, following exploration of the linear track, participation rates were elevated for both deep and superficial neurons, consistent with an experience-driven increase in overall hippocampal network activity^58–60^ (Figure 4 D). Importantly, the experience-dependent increase in participation was significantly stronger for superficial neurons than for deep neurons (Figure 4 D).

**Figure 4.**
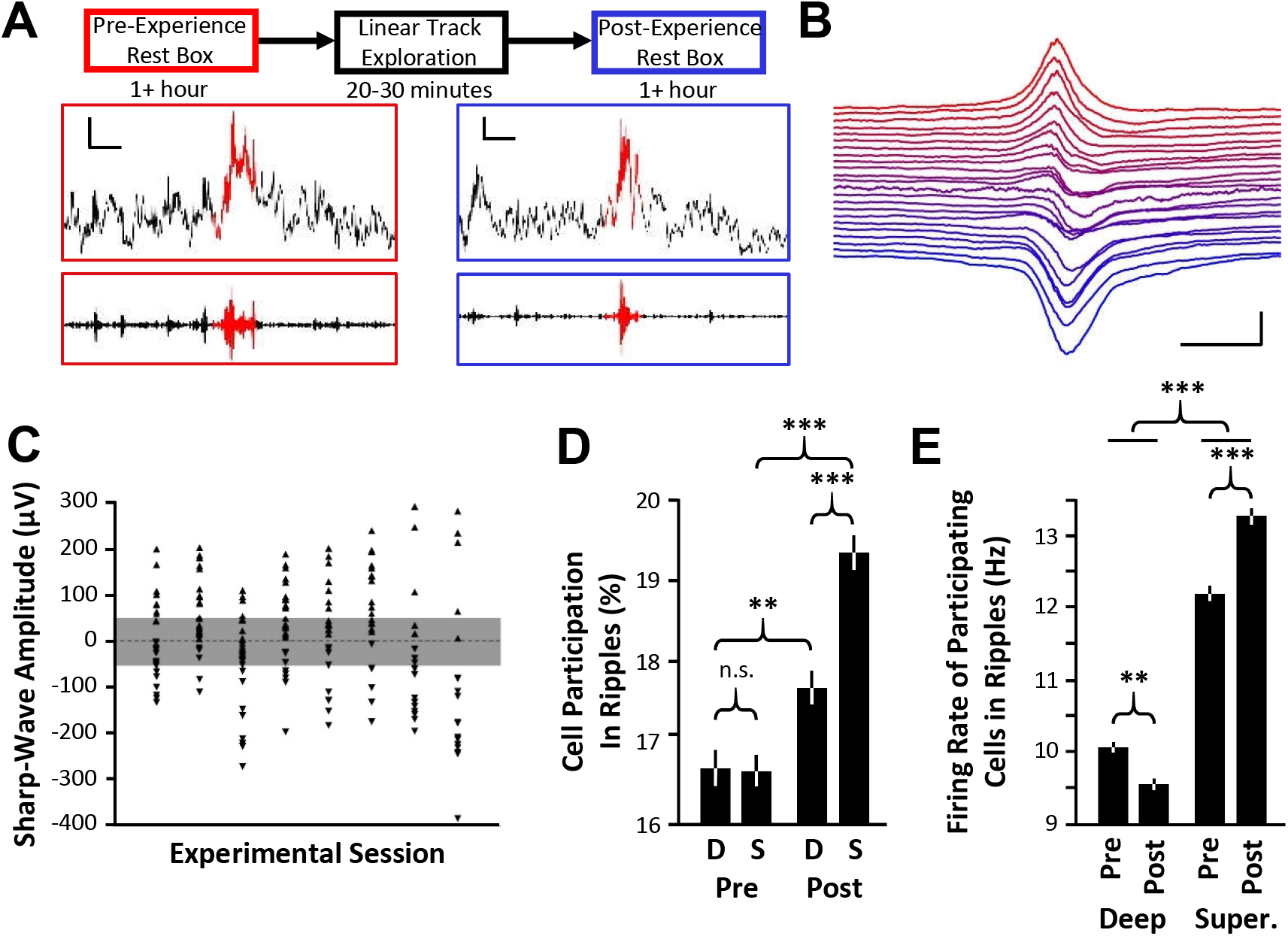
Deep and superficial pyramidal cell activity in SWRs is differentially affected by experience. **A**. Timeline of *in vivo* electrophysiology recording (top) and example SWRs recorded in the deep pyramidal layer (raw LFP middle, ripple-filtered LFP bottom, identified ripple in red) from pre-experience (left) and post-experience (right) phases. Scale bar = 0.2 s, 0.2 mV. **B**. Ripple-triggered average LFP trace (across 1,062 ripples) for each tetrode from one representative experimental session demonstrating range of recording depths. Tetrodes ordered by amplitude of sharp-wave deflection. Each tetrode baseline offset by 50 μV for visualization. Only tetrodes that recorded pyramidal neurons shown. Scale bar: 100 ms, 250 μV. **C**. For each experimental session, distribution of sharp-wave deflection amplitudes on each tetrode. Only tetrodes that recorded putative pyramidal neurons plotted. Tetrodes above or below the shaded region were classified as targeting deep or superficial layers, respectively. **D**. Mean ± S.E.M. percent of deep (D) or superficial (S) cells that participated per ripple in pre-experience (Pre) or post-experience (Post) rest sessions. **E**. For participating cells, mean ± S.E.M. firing rate for deep and superficial cells per ripple in pre- and post-experience rest sessions. Statistical tests and results: (D) Two-way ANOVA Deep vs. Superficial F(1,24948)=71.62, *p* < 10^−4^; Pre vs. Post F(1,24948)=14.36, *p* = 0.0002; Interaction F(1,24948)=15.25, *p* < 10^−4^. Multiple comparisons (Tukey HSD): Pre-Deep vs. Pre-Superficial *p* > 0.9; Pre-Deep vs. Post-Deep *p* = 0.007; Pre-Superficial vs. Post-Deep *p* = 0.005; All other comparisons *p* < 10^−4^. (E) Two-way ANOVA Deep vs. Superficial F(1,19539)=6.089, *p* = 0.014; Pre vs. Post F(1,19539)=714.6, *p* < 10^−4^; Interaction F(1,19539)=52.29, *p* < 10^−4^. Multiple comparisons (Tukey HSD): Pre-Deep vs. Post-Deep *p* = 0.0032; All other comparisons *p* < 10^−4^.

To more directly assess the impact of synaptic changes between CA3 and CA1 neurons independent of overall network activity levels, we next quantified the firing rates of individual deep and superficial neurons within SWRs. We hypothesized that synaptic depression from CA3 onto deep cells should reduce their firing rates in SWRs, while potentiation onto superficial cells should elevate their firing rates. In pre-experience SWRs, we observed a basal difference in the firing rates of deep and superficial neurons, consistent with the previously described impact of local inhibitory circuitry^41,61^. Deep CA1 neurons significantly reduced their firing rate from pre- to post-experience SWRs, while superficial neurons significantly increased their firing rate, consistent with opposite changes in their synaptic drive from area CA3 (Figure 4 E).

## DISCUSSION

We investigated the synaptic consequences of *in vivo* activity on hippocampal place cells and the effects of these changes on network reactivation in SWRs. Current models of memory formation argue that neural activity produced during active behavior generates long-lasting synaptic changes, which allow specific neural ensembles that represent prior experience to be reactivated during subsequent memory retrieval^2,62,63^. The expression of temporally compressed replay across the hippocampal network during SWRs is an established example of memory reactivation, and considerable work has sought to understand the cellular and synaptic mechanisms which underlie SWR-based replay. However, it is technically challenging to quantify the immediate consequences of *in vivo* activity on hippocampal synaptic function and the impact of such changes on network activity. Here, we address this issue by using CaMPARI2^11,12^ to permanently label active hippocampal neurons with sub-second temporal precision as animals traverse a novel linear track, followed by detailed synaptic analyses of active and inactive populations. We observe opposite synaptic changes following *in vivo* activity depending on whether the neurons reside in the deep or superficial sublayer of *stratum pyramidale*. Recently active deep pyramidal neurons display synaptic depression of CA3 inputs, while recently active superficial neurons are more potentiated at CA3 synapses than their inactive counterparts. These changes are reflected in the relative activity of these two groups during CA3-driven SWRs following a salient experience. Together with prior work^41,61^, our findings suggest a network model in which activity during exploration drives synaptic potentiation at CA3-to-superficial-CA1 synapses, resulting in elevated rates of superficial cell participation and firing rates in post-experience SWRs, while *in vivo* activity depresses CA3 input to deep pyramidal neurons, resulting in a reduction of firing rate and a relative suppression of participation of this cell population in SWRs.

A critical consideration for the interpretation of our findings is whether the differences observed in synaptic strength between photoconverted and unconverted neurons existed prior to the experience^30,31,64^ or whether they were produced as a consequence of *in vivo* activity during exploration^6,8,30,33,34,65^. Because we are unable to perform whole cell electrophysiology on the same neurons before and after exploration, we acknowledge that the experiments described here cannot definitively establish a causal link between *in vivo* activity and synaptic plasticity. However, several arguments support our interpretation that the synaptic differences we observe do not represent pre-existing structure, but are instead generated dynamically by *in vivo* activity. If the synaptic depression observed for synapses onto photoconverted deep pyramidal neurons (Figure 3 A-B) existed prior to the experience, that neural population would be expected to have reduced overall activity during the experience due to their weaker synaptic inputs, and thus would be less likely to be active at the time of photostimulation and consequently less likely to be photoconverted. Therefore, the strong correlation between photoconversion rates and weaker synapses onto deep neurons (Figure 3 A-B) indicate that these changes represent a consequence of *in vivo* activity rather than a pre-existing phenotype. In addition, we tracked superficial and deep pyramidal neurons from pre-experience to post-experience rest periods via extracellular *in vivo* electrophysiological tetrode recordings. In these recordings, both superficial and deep neurons show changes in SWR participation and firing rate from pre- to post-experience states which are entirely consistent with the differences we observe in synaptic strength between photoconverted and unconverted neurons. Indeed, if the increased synaptic strength onto photoconverted superficial neurons represented a pre-existing state, it is difficult to explain the significant changes we observe in the firing rates of individual superficial neurons from pre- to post-experience SWRs. Finally, considerable evidence supports a role for *in vivo* synaptic plasticity in the formation of place fields during active navigation and the establishment of stable spatial representation across the hippocampal network^6–9,20–22,66^, arguing that experience can induce synaptic plasticity *in vivo*. Thus, while we leave open the possibility that the synaptic differences we observe between photoconverted and unconverted neurons reflect pre-existing phenotypes, we interpret these findings as plasticity-induced changes driven by *in vivo* activity during experience.

Decades of work support the hypothesis that synaptic plasticity across the hippocampal network during exploration strongly influences the content of post-experience SWR-based replay, although the specific changes induced during experience to support off-line reactivation were previously unknown. Place cell ensemble activity during exploration is more strongly correlated with post- than pre-experience SWR content^34,58,67,68^, and this change can be abolished by inhibiting NMDAR function during behavior^32,33^. In addition, artificial stimulation of *ex vivo* hippocampal slices using physiological activity patterns recorded from freely behaving rats induces synaptic plasticity at CA3-to-CA1 connections, an effect which requires high levels of acetylcholine consistent with the strong cholinergic tone observed during active exploration^69^. Our results provide novel insight into the location and direction of *in vivo* synaptic plasticity which arises within the intact hippocampal network during experience, and further establish a link between these synaptic changes and the content of SWRs. Our work also clarifies the findings of several prior reports. Previous studies have identified a subset of hippocampal pyramidal cells which rigidly participate in both pre- and post-experience SWRs and a second, plastic subset which are selectively incorporated into post-experience SWRs following exploration of a maze^30,70^. While these studies did not identify the sublayers in which the rigid or plastic cells resided, our data indicate that superficial neurons are more likely to make up the plastic population, as potentiation of inputs to superficial cells selectively elevates their participation in post-experience SWRs. Consistent with this interpretation, rigid cells were reported to be higher firing and more likely to fire in a burst than plastic cells^30^, consistent with increased burstiness reported for deep neurons^38^. Furthermore, population-level quantification of synaptic weights before and after a novel experience via *in vivo* local field potential recordings reported a mixture of potentiation and depression^65^. As we observe a combination of potentiation and depression on superficial and deep pyramidal neurons, respectively, our findings help explain this variability.

To faithfully capture potential synaptic changes produced by *in vivo* activity, we sought to perform whole cell recordings as immediately as possible following the experience. Thus, the divergence we observe between photoconverted and unconverted neurons likely represents relatively rapid synaptic changes, and it is unclear from our data how long such changes may persist. The synaptic differences directly correlate to experience-dependent changes in neural activity during SWRs, arguing that plasticity during experience, particularly at CA3-to-CA1 synapses, plays a key role in determining neural activity patterns during SWR-based replay. Given that replay of recent experience can persist for many hours^33,71^, the experience-induced plastic changes we report here may represent long-term alterations to the synaptic architecture of the hippocampal network. In addition, while we have photolabeled neurons during active exploration, it is likely that the neurons which were photoconverted during periods of movement also participated in SWR-based replay during brief pauses on the linear track^24,25^. Several studies have established that the activity patterns arising during SWRs can produce plastic changes in synaptic or network function^72–74^, and it is possible that neural activity during on-task SWRs, rather than activity during navigation, are responsible for the potentiation/depression that we observe.

Our data identify bidirectional plasticity on two sub-populations of CA1 pyramidal neurons and have implications for the mechanisms of cell selection during replay. Previous work has identified a difference between superficial and deep pyramidal neurons in their connectivity to local inhibitory neurons which biases SWR participation toward superficial neurons^41^. Our data argue that this pre-existing bias is likely strengthened following experience-dependent potentiation of CA3 inputs to superficial neurons and depression of inputs to deep neurons. It is unclear what underlies the differences we observe in plasticity between deep and superficial neurons. It is known that superficial and deep neurons have unique gene expression patterns which may underlie differential function of metabotropic pathways^37,75^. Given the critical importance of metabotropic acetylcholine receptors for both long-term potentiation and depression^69,76^, we hypothesize that neuromodulatory cascades are likely different in deep vs. superficial neurons, possibly resulting in bidirectional plastic changes^77^. In addition, the synaptic changes we observe are measured via stimulation of a large population of Schaffer collateral axons, and may mask distinct, synapse-specific plasticity. Importantly, however, the strong correlation of *ex vivo* synaptic changes and *in vivo* activity patterns argues that our measurements faithfully capture meaningful alterations to the hippocampal network.

Superficial and deep cells receive differential input from the lateral and medial entorhinal cortex, two extra-hippocampal regions that convey distinct information content^39^. Accordingly, deep and superficial place cells encode experience differently depending on the presence or absence of reward^44^ or the complexity of the environment^48^. In our experimental paradigm, we utilized a relatively simple linear track and labeled neurons active only during periods of active movement across the track, which largely excluded labeling of neurons which encoded the reward areas at the ends of the track. Therefore, it is possible that under different behavioral paradigms or labeling strategies that a distinct population of deep and superficial cells would be labeled and a different population of synapses would be activated which may engage in unique forms of plasticity^78^.

## STAR METHODS

### Generation of AAV vector and virus

The pAAV-DJ plasmid was a gift from Dr. Wei Xu’s lab (UTSW). The CaMPARI2 gene was removed from pAAV-Syn-CaMPARI2 (Addgene #101060) and ligated into AAVdj-CaMKII-GFP backbone (Addgene #64545), replacing GFP, resulting in CaMPARI2 expression via the CaMKII promoter. The plasmid products were sequence verified before use in viral preparations using the following primers: CaMPARI2 forward 5′-TAGTTCTGGGGGCAGCGGGGGCCACCATGCTGCAGAAC-3′ and reverse 5′-CCAGAGGTTGATTATCGATATTACGTACGCAGGTCCTC-3′. Viral stock was produced via the Baylor University viral core facility.

### Viral injection, fiber implantation, and behavior

All CaMPARI2 experiments were performed according to procedures approved by the UT Southwestern Institutional Animal Care and Use Committee and Use Committee and followed US National Institutes of Health animal use guidelines. Male and female Long-Evans rats (350-500 g) aged 3-6 months were used. Animals were handled daily and food-restricted to 85–90% of their free-feeding weight and then trained to traverse a 1.8 m linear track to receive a sweetened milk reward (100 μl) at either end. Rats were trained for 20 minutes once per day for a total of 6 training days. Linear track training occurred in a room separate and visually distinct from the photolabeling room. After training, rats received bilateral viral injections, followed by a 14-day recovery period to allow for viral expression, followed by surgical implantation of optic fibers. Viral injections were performed following isoflurane anesthesia. Craniotomies were produced and three 600 nL injections of virus were made bilaterally at coordinates (relative to bregma): AP 3.9, 4.0, 4.1, ML ± 2.6, 2.8, 3.0, DV 2.5, 2.6, 2.7. Optic fiber pairs (400 μm diameter) housed in a lightweight plastic sheath were implanted bilaterally following isoflurane anesthesia at coordinates AP 4.0, ML ± 2.8, DV 2.3 and secured to skull screws via dental cement. On the experimental day, animals in the novel exploration cohort were placed on a 1.8-meter-long linear track in a room distinct from the training room, where they performed the trained task (running back and forth for liquid reward). 405 nm light (Thorlabs) was delivered through the implanted optic fibers only during periods of active movement across the track at 100% power, 3.7 mW, at 0.5 s pulses until 300 s of total labeling time was achieved, typically requiring 30-40 minutes of total exploration. Animals in the home cage cohort received 300 total seconds of identical light stimulation, without regard for movement or sleep/wake state. For both two-photon images and slice electrophysiology experiments, rats were deeply anesthetized with isoflurane immediately after photolabeling and rapidly decapitated within minutes of the final light stimulation pulse. Following surgical hippocampal isolation, acute sagittal 230-250 μm brain slices were cut in 4°C dissection buffer. Slicing was typically completed less than 30 minutes following the final light stimulation pulse. Individual slices were incubated in a recovery chamber saturated with 95% O_2_/5% CO_2_ at 34°C for 20 min and then kept at 24°C for a minimum of 45 min in artificial cerebrospinal fluid (ACSF) prior to imaging or electrophysiology.

### Imaging

Images were acquired on an Olympus Ultima IV Bruker with a 20X Olympus immersion objective, exciting at 920 nm or 1020 nm for native or photoconverted CaMPARI2, respectively. Each slice was imaged multiple times across the CA1 proximal-distal axis, obtaining images both inside and outside the fiber optic light cone and establishing a per-slice normalization value, as the baseline expression and signal strength was variable. Images were obtained from 18 slices across six animals, three in the novel exploration and three in the home cage cohort. The pyramidal cell location data was obtained by drawing a line perpendicular to the estimated border between *stratum radiatum* and *stratum pyramidale* and the center of the soma. Deep and superficial cells were identified as neurons with the center of their soma farther or nearer, respectively, 40 μm from the *stratum radiatum*/*stratum pyramidale* border. Cells were excluded from image analysis if cell somas were located 100 μm or more below the *stratum radiatum*/*stratum pyramidale* border (within *stratum radiatum*) or if observed to have a round or horizontally elongated somatic morphology. Cells were considered to be in the optic cone if they were within 150 μm of the vertical edge of the optic fiber.

### Slice electrophysiology

Cells were visualized by epifluorescence imaging using a water immersion objective (40X, 0.8 numerical aperture) on an upright Olympus BX51 WI microscope, with video-assisted infrared charge-coupled device camera (QImaging Rolera). During recordings, single slices were constantly perfused in a submersion chamber with 30°C oxygenated ACSF with the addition of 50 μM picrotoxin. Patch pipettes were pulled to a final resistance of 3 to 5 MΩ from filamented borosilicate glass on a Sutter P-1000 horizontal puller. Superficial CA1 pyramidal cell somata were defined as cell bodies approximately 0-40 μm inside the pyramidal layer from the border of *stratum pyramidale* and *stratum radiatum*. Deep CA1 pyramidal cell somata were identified as approximately 50 μm or more from the *stratum pyramidale*/*stratum radiatum* border. Data were low-pass–filtered at 10 kHz and acquired at 2 kHz with an Axon MultiClamp 700B amplifier and an Axon Digidata 1550B data acquisition system under the control of Clampex 10.6 (Molecular Devices). Synaptic currents were evoked by electrical stimulation with an isolation unit through a glass stimulation monopolar electrode filled with ACSF and placed in the *stratum radiatum* (1 ms, 0.2-0.7 mA, 0.1 Hz). Synaptic responses were monitored at different stimulation intensities before establishing baseline recording at approximately 50% of the maximal response. For paired-pulse ratio measurements, synaptic responses were obtained while holding the cell at −70 mV and delivering paired stimulations at 20 Hz. The paired-pulse ratio is calculated as the amplitude of the second response divided by the amplitude of the first. To measure the AMPA/NMDA ratio, the membrane voltage was held at +40 mV. Responses to *stratum radiatum* stimulation included both AMPAR- and NMDAR-driven currents. AP5 (d,l-2-amino-5-phosphonovaleric acid, 100 mM) was bath applied and AMPAR-isolated currents were obtained using identical stimulation. The NMDA component was obtained by digital subtraction of the ACSF-only current at +40 mV and the residual current after AP5 application. Access resistance (10 to 30 MΩ) was monitored throughout the experiment and cells where it changed more than 20% were discarded from further analysis. A total of 14 rats were used for slice electrophysiology.

### *In vivo* electrophysiology

The dataset used in this study has been previously analyzed^79^. All *in vivo* electrophysiology experiments were performed according to the procedures approved by the Johns Hopkins University Animal Care and Use Committee and followed US National Institutes of Health animal use guidelines. A detailed description of the behavioral training and electrophysiological recording has been described previously^25^ and is summarized here. Following training, adult male Long-Evans rats were implanted with a microdrive array (25-30 g) containing 40 independently adjustable, gold-plated tetrodes aimed at CA1 of dorsal hippocampus. Recordings of spike data (32 kHz, 600-6000 Hz bandpass) and local field potentials (LFP, 3.2 kHz, 0.1-500 Hz bandpass) were obtained during free exploration of a linear track for reward at either end and during one hour rest periods immediately before and after linear track exploration in which the rat was confined to a paper-lined, 30-cm-diameter ceramic plate atop an inverted vase within a 60-cm x 60-cm room with opaque walls (the rest box). Only tetrodes in which single units were identified were included in subsequent analyses. Post-experiment lesion histology^79^ confirmed recording locations in the CA1 pyramidal area but were incapable of determining precise deep vs. superficial targeting due to the size of the lesions. To identify sharp-wave/ripples (SWRs), the LFP was bandpass filtered between 150 and 250 Hz, and the absolute value of the Hilbert transform of this filtered signal was smoothed (Gaussian kernel, SD 12.5 ms). This processed signal was averaged across all tetrodes and SWRs were identified as local peaks with an amplitude >3 SD above the mean. The start and end boundaries of each SWR were defined as the point when the signal crossed the mean. For each tetrode, the sharp-wave deflection was quantified as the mean amplitude across all SWRs of the 5-15 Hz bandpass filtered signal at maximum ripple power. Tetrodes with a sharp-wave deflection greater than 50 μV were classified as targeting the deep sublayer; tetrodes with a sharp-wave deflection less than −50 μV were classified as targeting the superficial sublayer. Only units classified as putative excitatory neurons (based on spike waveform and mean firing rate^25^) with mean spike amplitude of 200 μV or greater on at least one electrode of the tetrode were included in analyses.

### Data and statistical analysis

ImageJ software was used to quantify the conversation ratios and radial location of CA1 pyramidal cells. Clampfit (Molecular Device) was used for the analysis of slice electrophysiological data and trace extraction. All data were analyzed in Prism 9.0 (GraphPad) or MATLAB 2022a (MathWorks). Statistics for each test are included in figure legends. All data and analysis code are freely available upon request.

### Solutions

Dissection buffer for imaging: NaHCO3 (25 mM), KCl (2.5 mM), NaH2PO4 (1.25 mM), MgCl2 (7 mM), CaCl2 (0.75 mM), sucrose (211 mM), D-glucose (11 mM), and Na-pyruvate (3 mM). ACSF for imaging: NaCl (125 mM), NaHCO3 (26 mM), KCl (2.5 mM), NaH2PO4 (1.25 mM), MgCl2 (1 mM), CaCl2 (2 mM), D-glucose (11 mM). ACSF pH was approximately 7.3 after oxygenation with 95%/5% O2/CO2. Dissection buffer for electrophysiology: 225 mM sucrose, 3 mM KCl, 1.25 mM NaH2PO4, 26 mM NaHCO3, 10 mM d-(+)-glucose, 7 mM MgSO4, 0.5 mM CaCl2, and 2 mM kynurenic acid. ACSF for electrophysiology: 126 mM NaCl, 3 mM KCl, 1.25 mM NaH2PO4, 26 mM NaHCO3, 10 mM d-(+)-glucose, 1.3 mM MgSO4, and 2.5 mM CaCl2. Internal solution: 120 CsMeSO3, 10 CsCl, 10 HEPES, 10 EGTA, 5 Na-phosphocreatine, 4 ATP (Mg2+ salt), 0.4 GTP (Na+ salt), pH adjusted to 7.3 with CsOH, osmolarity adjusted to 300 ± 10 mOsm.

## Acknowledgements

We thank Dr. Wei Xu, Dr. Xin Shao, and Harshida Pancholi for sharing reagents and technical expertise. This work was supported by National Institutes of Health: R01MH117149 (LJV), R01NS104829 (BEP), and R01NS108424 (TFR).

